# Bacterial Genome wide association studies (bGWAS) and transcriptomics identifies cryptic antimicrobial resistance mechanisms in *Acinetobacter baumannii*

**DOI:** 10.1101/864462

**Authors:** Chandler Roe, Charles H.D. Williamson, Adam J. Vazquez, Kristen Kyger, Michael Valentine, Jolene R. Bowers, Paul D. Phillips, Veronica Harrison, Elizabeth Driebe, David M. Engelthaler, Jason W. Sahl

## Abstract

Antimicrobial resistance (AMR) in the nosocomial pathogen, *Acinetobacter baumannii*, is becoming a serious public health threat. While some mechanisms of AMR have been reported, understanding novel mechanisms of resistance is critical for identifying emerging resistance. One of the first steps in identifying novel AMR mechanisms is performing genotype/phenotype association studies. However, performing genotype/phenotype association studies is complicated by the plastic nature of the *A. baumannii* pan-genome. In this study, we compared the antibiograms of 12 antimicrobials associated with multiple drug families for 84 *A. baumannii* isolates, many isolated in Arizona, USA. *in silico* screening of these genomes for known AMR mechanisms failed to identify clear correlations for most drugs. We then performed a genome wide association study (GWAS) looking for associations between all possible 21-mers; this approach generally failed to identify mechanisms that explained the resistance phenotype. In order to decrease the genomic noise associated with population stratification, we compared four phylogenetically-related pairs of isolates with differing susceptibility profiles. RNA-Sequencing (RNA-Seq) was performed on paired isolates and differentially expressed genes were identified. In these isolate pairs, we identified four different potential mechanisms, highlighting the difficulty of broad AMR surveillance in this species. To verify and validate differential expression, amplicon sequencing was performed. These results suggest that a diagnostic platform based on gene expression rather than genomics alone may be beneficial in certain surveillance efforts. The implementation of such advanced diagnostics coupled with increased AMR surveillance will potentially improve *A. baumannii* infection treatment and patient outcomes.

## Introduction

Antimicrobial resistance (AMR) has the potential to become a global health emergency and is expected to kill more people than cancer by the year 2050 (1). Multidrug resistance in *Acinetobacter baumannii* is now recognized as a major public health concern, resulting in the World Health Organization (WHO) declaring *A. baumannii* a priority 1 pathogen (2). *A. baumannii* is primarily a nosocomial pathogen (3) that affects immunocompromised patients, causing a variety of afflictions including pneumonia, septicemia, meningitis, and death (4, 5). Treatment of *A. baumannii* infections has become increasingly difficult due to the emergence of multidrug resistance; pan-resistant *A. baumannii* strains (6–8), including strains resistant to last-resort drugs such as colistin (9), have been identified in Asia and Europe.

Known mechanisms that confer AMR in *A. baumannii* include penicillin binding proteins (10), enzymes (11), porin defects (12), DNA methylation (13), and efflux pumps (14). Efflux pumps that confer resistance in *Acinetobacter* are classified into four families: multidrug and toxic compound extrusion (MATE), resistance–nodulation–division (RND) family, major facilitator superfamily (MFS), and small multidrug resistance (SMR) (15). Additionally, mutations in promoter regions can lead to overexpression of some efflux systems, including AdeFGH (14), which has been shown to lead to resistance to multiple antimicrobial families.

Resistance mechanisms have also been reported for specific drug families used to treat *A. baumannii* infections. Perhaps the most studied family is beta-lactams, including carbapenems (e.g. meropenem and imipenem), which are used to treat nosocomial infections (16, 17). Carbapenem resistance has been associated with the action of carbapenem-hydrolyzing class D beta-lactamases (CHDLs), including bla_OXA-23_, bla_OXA-24_, and bla_OXA-58_ (18). The *ampC* cephalosporinase is a class C beta-lactamase that is broadly conserved across *A. baumannii A.* (19) and has been associated with resistance to narrow spectrum cephalosporins (20). Additionally, bla_OXA-51-like_ genes are highly conserved across the *A. baumannii* species, as well as other *Acinetobacter spp*. (21); these genes confer resistance to carbapenems when in close proximity to insertion element ISAba1 (22).

Aminoglycoside resistance in *A. baumannii* has been associated with the actions of aminoglycoside modifying enzymes (AMEs) including *aacC1*, *aphA6*, *aadA1*, and *aadB* (23), the 16S rRNA methyltransferase *armA* (24), as well as through efflux action of AdeABC and AbeM, although the efflux effect was limited (23). Resistance to macrolides in *A. baumannii* has primarily been associated with target site alteration in Dfr (25) encoded by *folA*, the presence and activity of the *tetM* gene (26), and through the action of efflux pumps (27). Finally, quinolone resistance in *A. baumannii* has been linked to quinolone resistance-determining regions (QRDRs) (28), including mutations in *parC*, *gyrA,* and *gyrB*. Specifically, the *gyrA* S82L mutation has previously been shown to confer resistance to quinolones (29); two separate mutations, S83L and G80V, have also been demonstrated to confer quinolone resistance in *A. baumannii* (30).

In recent years, multiple databases have been developed and maintained that include genomic regions associated with antimicrobial resistance. These databases include CARD (31), ResFinder (32), ARG-ANNOT (33), ARDB (34), and MEGARes (35). To identify potential resistance mechanisms, genomes are screened against these databases and if genes associated with resistance are identified and conserved, then resistance patterns are inferred (36). However, genomics doesn’t capture expression profiles, including gene expression induction, which prevents accurate genotype to phenotype associations in some organisms (37).

Current treatment regimens for *A. baumannii* infections start with broad spectrum cephalosporins such as ceftazidime or cefepime, or a carbapenem (e.g. imipenem) (38). For drug resistant pathogens, polymyxins such as colistin are used, although emerging resistance has been reported (39) and the treatment can be toxic (40). Other drugs, including tigecycline (41) and minocycline (42) have been used to treat resistant strains, although resistance to these therapies has also been observed, prompting research into combination therapies (43) that overcome these limitations. However, pan-resistance in *A. baumannii* (44) has the potential to undermine all current treatment regimens and necessitates a better understanding of genotype/phenotype associations for improved surveillance efforts and targeted therapy.

Research into genotype/phenotype associations in *A. baumannii* is complicated by the highly plastic nature of the pan-genome (45). One example of this phenomenon is the biofilm associated protein (Bap) (46, 47), which, based on an *in silico* screen of more than 117 complete *A. baumannii* genomes, is conserved in only 2 genomes (unpublished). This demonstrates that mechanisms associated with a phenotype may not be broadly distributed across diverse isolates of this species. While true for virulence, similar patterns exist for AMR genes that are variably conserved within a highly plastic species (21).

In this study, we analyzed over 100 *Acinetobacter* isolates, largely isolated in Arizona, USA, in an effort to identify broadly conserved as well as cryptic mechanisms of AMR. Implementing an iterative approach, we searched common AMR gene databases for known mechanisms, performed a genome wide association study (GWAS) to identify potentially new mechanisms, and performed RNA-Sequencing (RNA-Seq) to compare gene expression profiles between isolates with variable AMR phenotypes. The results provide additional detail to understand AMR mechanisms in *A. baumannii* and identify targets for advanced diagnostics that will provide appropriate therapies for more effective patient treatment.

## Methods

### Isolate description and growth

A total of 107, largely geographically confined isolates, were identified for sequencing based on collection from different body sites and clinical matrices. All isolates were classified as *A. baumannii* based on orthogonal, clinical laboratory techniques. A description of all sequenced isolates is shown in Table S1. Samples were streaked from glycerol stocks onto Mueller Hinton (MH) (Hardy Diagnostics, Santa Maria, CA) agar plates and incubated at 37°C for 24 h. Inoculated plates were checked for appropriate colony growth and morphology the following day prior to DNA extraction.

**Table 1:** A list of antimicrobials screened in the current study

| Antimicrobial | Abbreviation | Family | Publication | Susceptible breakpoint | Resistant breakpoint | Reference |
| --- | --- | --- | --- | --- | --- | --- |
| cefepime | PM | B-lactam | Endimiani et al. 2008 | $\geq 32$ | $\leq 8$ | CLSI |
| cefuroxime | XM | B-lactam | Ahmed et al. 2012 | $\geq 128$ | $\leq 32$ | N/A |
| gentamicin | GM | aminoglycoside | Hamidian et al. 2012 | $\geq 16$ | $\leq 4$ | CLSI |
| ceftazidime | TZ | B-lactam | Lee et al. 2006 | $\geq 16$ | $\leq 4$ | CLSI |
| trimethoprim | TR | pyrimidine inhibitor | McCracken et al. 2009 | $\geq 32$ | $\leq 4$ | EUCAST<sup>1</sup> |
| azithromycin | AZ | macrolide | Fernandez Cuenca et al. 2003 | $\geq 256$ | $\leq 8$ | N/A |
| ceftriaxone | TX | B-lactam | Bush et al. 1995 | $\geq 64$ | $\leq 8$ | CLSI |
| aztreonam | AT | B-lactam | Xia et al. 2014 | $\geq 32$ | $\leq 8$ | CLSI |
| erythromycin | EM | macrolide | Damier-Piolle et al. 2008 | $\geq 8$ | $\leq 0.5$ | CLSI<sup>2</sup> |
| piperacillin | PP | B-lactam | Shi et al. 1996 | $\geq 128$ | $\leq 16$ | CLSI |
| levofloxacin | LE | fluroquinolone | Lee et al. 2006 | $> 1$ | $\leq 0.5$ | EUCAST |
| ciprofloxacin | CI | fluroquinolone | Chiu et al. 2010 | $\geq 4$ | $\leq 1$ | CLSI |

<sup>1</sup>Enterobacteriaceae
<sup>2</sup>Enterococcus

### Genomic DNA extraction and Sequencing

Genomic DNA was extracted from a single isolated colony for each sample using the DNeasy Blood and Tissue Kit (Qiagen, Valencia, CA, USA) following the recommended protocol for Gram-negative bacteria. Sample DNA was fragmented using the QSonica q800 ultrasonic liquid processor (QSonica, Newtown, CT, USA). Sonication parameters were optimized to produce fragment sizes of 600 to 700 base pairs (time: 3 minutes, pulse: 15s (Pulse On), 15s (Pulse Off), amplitude: 20%). Libraries were size selected using Agencourt AMPure XP beads (Beckman Coulter, Brea, CA) in order to remove small and large fragments outside of the required size range. Genome libraries were prepared using the KAPA Hyper Library Preparation Kit with Standard PCR Library Amplification (Kapa Biosystems, Wilmington, MA) and sequenced on an Illumina MiSeq using V3 sequencing chemistry (Illumina Inc., San Diego, CA).

For MinIon sequencing, DNA was extracted with the GenElute Bacterial Genomic DNA kit (Sigma-Aldrich Inc., St. Louis, MO), taking care to limit DNA shearing. Long read sequencing was performed using Oxford Nanopore technologies on a MK1B MinION device using a R9.4 flow cell. The DNA library was prepared using the SQK-LSK109 Ligation Sequencing kit in conjunction with the PCR-Free Native Barcode Expansion kit following manufacturer’s protocol (downloaded from https://nanoporetech.com/resource-centre/protocols/ on March 20, 2019) without the optional shearing steps to select for long reads.

### Sequence assembly and MLST typing

Illumina-derived whole genome sequence data was assembled with SPAdes v3.10 (48). Contigs that aligned against known contaminants or contained an anomalously low depth of coverage compared to the average depth of coverage on a per genome basis were manually removed. The MLST profiles were extracted from whole genome sequence (WGS) assemblies using BLAST-based methods (49) using both the Oxford (50) and Pasteur systems (51). Annotation on all genomes was performed with Prokka v1.13 (52). Hybrid assemblies were generated with combined Illumina and MinION data with Unicycler v0.4.8-beta (53). Assemblies were polished with Pilon v1.22 (54).

### Global phylogenetics of *Acinetobacter*

*Acinetobacter* genome assemblies were downloaded from GenBank on March 13th, 2018. All genome assemblies were aligned against the *A. baumannii* genome AB307-2094 (CP001172.1) with NUCmer v3.1 (55) in conjunction with NASP v1.1.2 (56). SNPs that fell within duplicated regions, based on a reference self-alignment with NUCmer, were filtered from downstream analyses. For rapid evaluation, an approximate maximum likelihood phylogeny was inferred on a concatenation of 1,523,968 single nucleotide polymorphisms (SNPs) with FastTree v2.1.8 (57); SNPs were retained if they were conserved in >90% of all genomes.

### Antimicrobial resistance profiling

Antimicrobial resistance phenotypic profiles were identified for cefepime (PM), cefuroxime (XM), gentamicin (GM), ceftazidime (TZ), trimethoprim (TR), azithromycin (AZ), ceftriazone (TX), aztreonam (AT), erythromycin (EM), piperacillin (PP), levofloxacin (LE), and ciprofloxacin (CI). A list of all drugs and resistance breakpoints used are shown in Table 1. Drugs were selected from published resistance patterns in the literature (58–68). Samples were streaked from glycerol stocks onto Mueller Hinton (MH) (Hardy Diagnostics, Santa Maria, CA) agar plates and incubated at 37°C for 24h. A single isolated colony was picked and inoculated into 10mL of MH broth. Liquid cultures were incubated with shaking at 37°C overnight. The following morning, 100µL of each overnight culture were transferred into 9.9mL of fresh MH broth. Cultures were incubated with shaking at 37°C until optical density (OD_600_) measurements reached 0.5-0.8, indicating log phase growth. 50µL of culture was inoculated onto new 15 x 150mm MH agar plates and spread uniformly across the medium with a sterile cell spreader. Six different antimicrobial E-test strips (bioMérieux, France) were applied to the surface of the agar as directed by the manufacturer. Plates were incubated at 37°C for 16-18hrs and minimum inhibitory concentrations (MIC) were determined by visual inspection following the recommended manufacturer guidelines. For paired isolates, MIC tests were performed on different days.

### *A. baumannii* phylogeny and isolate pairing

Once the set of *A. baumannii* were identified, a phylogeny was generated for confirmed genomes (n=84). Raw WGS data were aligned against AB307-0294 with BWA-MEM v0.7.7 (69) and single nucleotide polymorphisms (SNPs) were identified with the UnifiedGenotyper method in GATK v3.3.1 (70, 71). SNPs that fell into duplicate regions of the reference, based on a NUCmer self-alignment, were removed from downstream analyses. All SNP calling methods were wrapped by the NASP pipeline. A maximum likelihood phylogeny was inferred on a concatenation of 182,766 SNPs with IQ-TREE v1.6.1, using the TVMe+ASC+R5 model. Paired genomes were identified by low phylogenetic distance and variable antibiograms (Table S2).

**Table 2:** Paired isolate antimicrobial susceptibility

| Intermediate isolate | Resistant Isolate | Drug | Pair number | PubMLST/Pasteur | Resistant MIC | Other MIC |
| --- | --- | --- | --- | --- | --- | --- |
| TG22627 | TG22182 | TX,TZ | 1 | ST368/ST2 | >256, >256 | 48, 8 |
| TG31302 | TG31986 | PM | 2 | ST1961/ST78 | >256 | 12 |
| TG31307 | TG29392 | XM,TX | 3 | ST1961/ST78 | >256, >256 | 64, 32 |
| TG60155 | TG22653 | PM | 4 | ST208/ST2 | >256 | 16 |

### Global phylogenetic analysis

All *A. baumannii* genomes (n=3,218) were downloaded from the Assembly database in GenBank (72) on September 19th, 2018. Genomes were filtered if they: 1) contained greater than 200 ambiguous nucleotides (n=860); 2) contained greater than 400 contigs (n=189); 3) had a genome assembly size <3684234 or >4297137 (n=51), or; 4) had an average MASH (73) distance greater than 0.0252 (∼97.5% average nucleotide identity) (n=20). Genomes passing through all filters (n=2183) were aligned against *A. baumannii* AB307-2094 (6) with NASP in conjunction with NUCmer. A maximum likelihood phylogeny was inferred on a concatenation of 101,608 SNPs with IQ-TREE v1.6.1, using the TVM+F+ASC+R10 model, and rooted with an *A. nosocomialis* genome sequenced in this study (TG22170; RFEG00000000).

### Comparative analysis of paired isolate genomes

To identify coding region differences between paired isolates, the large-scale blast score ratio (LS-BSR) (74) tool was run on paired genomes in conjunction with BLAT (75). The order of genes between isolates was visualized with genoPlotR (76).

### Genome wide association studies (GWAS)

To identify genotype/phenotype associations, regions identified by LS-BSR were compared between resistant/susceptible phenotypes from each drug. Regions were first identified that had a blast score ratio (BSR) value (77) of >0.8 in one phenotype and a BSR value of <0.4 in the other phenotype. Correlations between groups was identified with a point biserial correlation method. In addition to differences in coding region sequences (CDSs), individual SNPs and indels were identified through the analysis of Kmers. In this approach, the reverse complement was taken for all genomes so that both strands were included in the analysis. All 21-mers were then identified with Ray-surveyor (78) and placed into a presence absence matrix; the choice of 21-mers was to ensure a short enough length to hopefully identify single mutations. The frequency of Kmers in each phenotype was then calculated with a custom Python script (https://gist.github.com/jasonsahl/e9516b2d940ad2474ba6e97f5b856440).

### Machine learning approach for AMR mechanism identification

Associations of Kmers of length 21 with each AMR mechanism were identified with Kover v2.0.0 (79) using default parameters. Annotation for Kmers was performed by mapping Kmers against annotated coding regions with BLASTN.

### *in silico* screen of antimicrobial resistance elements

To find previously characterized antimicrobial resistance mechanisms, we screened paired genomes with LS-BSR in conjunction with Diamond (80) and the Comprehensive Antimicrobial Resistance Database (CARD) (31). We also selected resistance genes from the literature associated with antimicrobials screened in this study (Table S3) using the same methods.

**Table 3:** Kover results for AMR across *A. baumannii*

|  | #Resistant<br>isolates | #Susceptible<br>isolates |  | Equivalent<br>rules |  |
| --- | --- | --- | --- | --- | --- |
| Drug |  |  | Importance |  | #loci |
| AT | 57 | 7 | 1 | 105 | 9 |
| TZ | 68 | 9 | 1 | 10000 | 1922 |
| PM | 61 | 13 | 1 | 665 | 37 |
| LE | 71 | 12 | 1 | 55 | 8 |
| GM | 57 | 16 | 0.87 | 1 | 1 |
| AZ | 50 | 6 | 1 | 893 | 81 |
| XM | 67 | 11 | 0.86 | 2193 | 225 |
| TR | 84 | 0 | N/A | N/A | N/A |
| TX | 67 | 3 | N/A | N/A | N/A |
| EM | 81 | 0 | N/A | N/A | N/A |
| PP | 69 | 5 | 1 | 10000 | 2309 |
| CI | 73 | 11 | 1 | 143 | 23 |

### Antimicrobial exposure and RNA extractions

Samples identified as paired isolates based on phylogenetic relatedness and differing antibiogram profiles were streaked for isolation from glycerol stocks onto MH agar plates and incubated overnight at 37°C. For each sample a single colony was picked and inoculated into 10mL of MH broth and incubated with shaking at 37°C overnight. The following morning 100µL of each culture was inoculated into 9.9mL of fresh media and OD_600nm_ was monitored until cultures reached log phase growth OD_600nm_ of approximately 0.5-0.8. 500µL of each sample, as well as susceptible control strain *Staphylococcus aureus subsp. aureus* (ATCC 29213), were aliquoted in triplicate into 2mL microcentrifuge tubes. Each sample was treated with sub-MIC concentrations of the designated antimicrobial, at one half of the previously recorded MIC value. Cultures were then incubated for 30min with shaking at 37°C. Two volumes of RNAprotect Bacteria Reagent (Qiagen, Valencia, CA, USA) were added to all samples and incubated at room temperature for 5min, followed by centrifugation for 10min at room temperature, at a speed of 5000 x g. The supernatant was decanted and the treated cell pellets were stored at −80°C. Total RNA was extracted using the RNeasy Mini Kit (Qiagen, Valencia, CA, USA) following recommended protocol #4 beginning at step 7 and continuing to protocol #7. A DNase I treatment was included for step 2 in protocol #7. Extracted RNA was immediately stored at −80°C.

### mRNA isolation

RNA quality and quantity were checked by Agilent 2100 Bioanalyzer with the RNA 6000 Nano Kit (Agilent Technologies, Santa Clara, CA, USA). mRNA was isolated from total RNA using the MICROBExpress kit (Thermo Fisher Scientific, Waltham, MA) following the manufacturer’s protocol. Isolated mRNA was quantified and checked for rRNA depletion on the bioanalyzer with an additional RNA Nano chip prior to sequencing.

### RNA-seq preparation, sequencing, assembly

Previously isolated mRNA was prepared for transcriptome sequencing using the TruSeq Stranded mRNA, HT kit (Illumina, San Diego, CA) following the High Sample (HS) protocol. Prepared samples were quantified and checked for quality, then pooled in equimolar concentrations. Library pools were loaded into an Illumina High Output NextSeq 2 x 150bp kit, according to manufacturer recommendations for sequencing on the Illumina NextSeq 550 platform. The transcriptomes were assembled with metaSPAdes (81) using default settings. For targeted amplicon studies, complementary DNA (cDNA) was generated with the SuperScript IV VILO RT-PCR Master Mix with ezDNase enzyme (Invitrogen, Carlsbad, California), following manufacturer’s recommendations.

### Differential expression (DE) analysis

For each isolate pair, coding and intergenic regions identified with LS-BSR and prodigal were combined for complete genomes, then dereplicated with USEARCH v10 at an ID of 0.98. RNA-Seq reads were aligned against these regions with BWA-MEM and read counts were called on the resulting BAM file with Salmon v0.13.1 (82). Differential expression (DE) analysis was performed with DESeq2 (83). The p-values were corrected using the Benjamini-Hochberg (84) correction.

### Amplicon sequencing (AmpSeq)

Polymerase chain reaction (PCR) primers were designed for differentially expressed regions identified in the RNA-Seq analysis (Table S4); a constitutively expressed target (locus tag: IX87_18340), based on analysis of RNA-Seq data, was included for normalization. cDNA was amplified with the following protocol: 1X Promega PCR Master Mix (Promega, Fitchburg, WI), 2.5µL cDNA template, and multiplexed primer concentrations are listed in Table S4. Gene specific PCR parameters were as follows: initial denaturation at 95°C for 2m, 30 cycles of denaturation at 95°C for 30s, annealing at 55°C for 30s, and extension at 72°C for 45s, with a final extension at 72°C for 5m. Included on each primer was a universal tail (85), which facilitated Illumina index ligation. Samples were indexed with the following final concentrations: 1X HiFi HotStart Readymix (Kapa Biosystems Inc., Wilmington, MA), 0.4µM of each indexing primer, and 2µL of gene specific PCR product. The indexing PCR parameters were as follows: initial denaturation at 98°C for 2m, 6 cycles of denaturation at 98°C for 30s, annealing at 60°C for 20s, and extension at 72°C for 30s, with a final extension at 72°C for 5m. Following each PCR, a 1X Agencourt AMPure bead (Beckman Coulter, Brea, CA) clean-up was performed according to manufacturer’s instructions. All amplicons were normalized with SequalPrep (Thermo Fisher Scientific, Applied Biosystems), pooled, and sequenced on the Illumina MiSeq platform (Illumina Inc., San Diego, CA).

**Table 4:** Associated genotype/phentotype genomic regions

| Drug | #Resistant<br>(R) | #Susceptible<br>(S) | #unique 21-mers<br>(R) | #unique 21-mers<br>(S) | #unique<br>genes (R) | #unique<br>genes (S) |
| --- | --- | --- | --- | --- | --- | --- |
| AT | 57 | 7 | 0 | 0 | 0 | 0 |
| AZ | 50 | 6 | 0 | 0 | 0 | 0 |
| CL | 73 | 11 | 0 | 0 | 0 | 0 |
| EM | 81 | 0 | N/A | N/A | N/A | N/A |
| GM | 57 | 16 | 0 | 0 | 0 | 0 |
| LE | 71 | 12 | 0 | 0 | 0 | 0 |
| PM | 61 | 13 | 0 | 0 | 0 | 0 |
| PP | 70 | 5 | 0 | 3 | 0 | 0 |
| TX | 67 | 3 | 3* | 47 | 1* | 0 |
| TZ | 68 | 9 | 0 | 0 | 0 | 0 |
| XM | 67 | 11 | 0 | 0 | 0 | 0 |

\*Present in n-1 genomes

### AmpSeq analysis

Raw AmpSeq data were aligned against predicted amplicons with Kallisto v0.45.0 (86). Counts were normalized based on the median read counts between all samples. The difference between the raw read counts of the target and the housekeeping gene was identified for each sample. The average delta was then identified for each set of resistant and intermediate genomes and the delta Ct was calculated. The average deltas were compared between resistant and intermediate samples and a p-value was calculated with a two-sided T-test.

### Data availability

All data were deposited to appropriate databases and linked under BioProject PRJNA497581. Links to specific samples are shown in Table S1.

## Results

### Identification of isolates analyzed in the current study

In this study, we sequenced 107 isolates identified by laboratory methods to be *A. baumannii*. These isolates were retrospectively identified from our collection and sequenced to reflect a range of years and isolation sources (Table S1). Of the 107 genomes sequenced in this study, only 84 were confirmed *A. baumannii* (Table S1) isolates based on a global WGS phylogenetic analysis (Figure 1). In order to define and add context to the phylogenetic diversity of genomes sequenced in this study, more than 3,000 publicly available *A. baumannii* genomes were included in the analysis (Figure S1).

**Figure 1:**
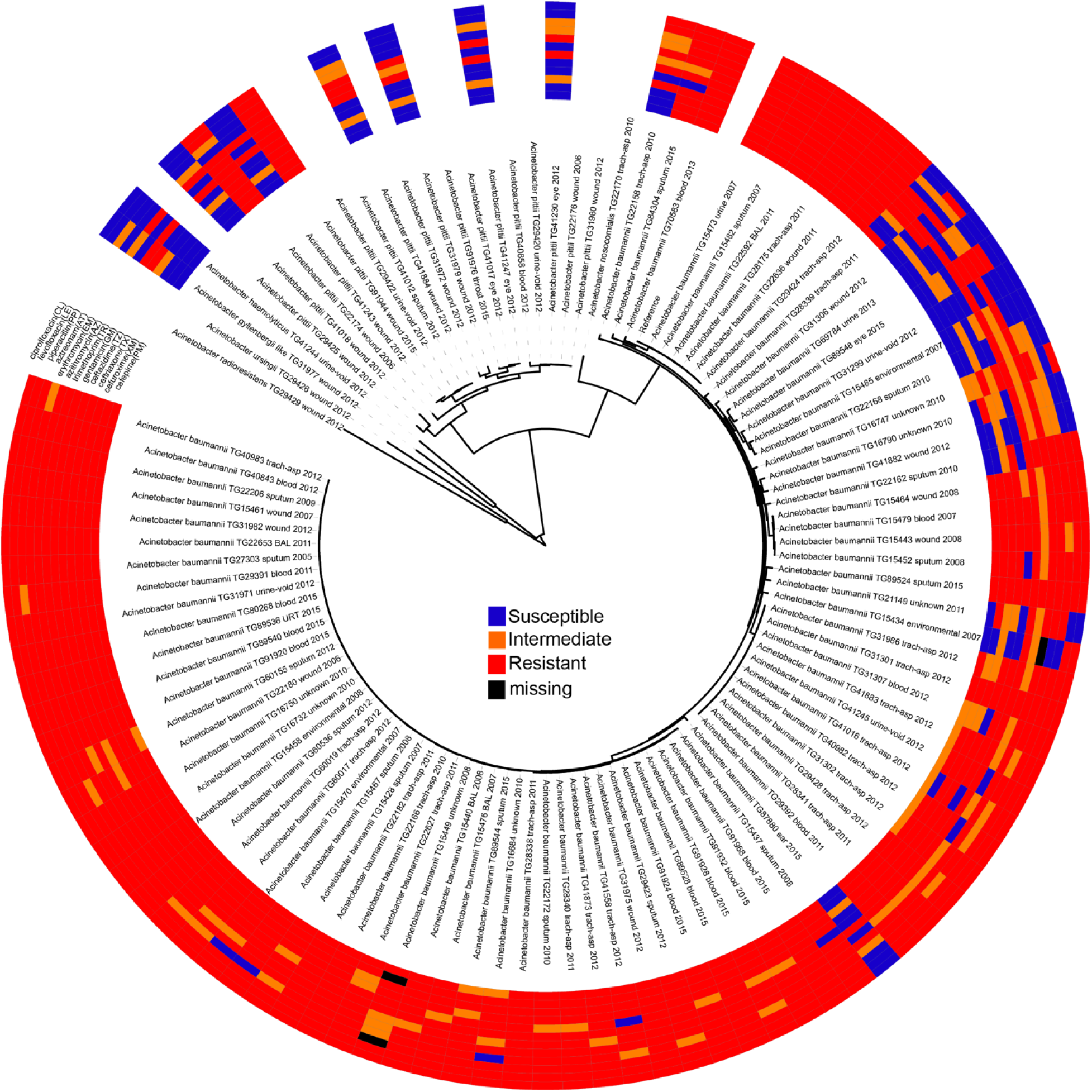
A maximum-likelihood phylogeny of genomes sequenced in this study based on a concatenation of 50,869 core genome SNPs. Each genome is annotated with its antimicrobial susceptibility profile across 12 drugs. The annotations were visualized with the Interactive tree of life (111).

### Antimicrobial resistance profiles of isolates analyzed

AMR profiles were identified for 95 of the isolates across 12 drugs, including all *A. baumannii* (Figure 1, Table S2). Twelve isolates were excluded due to either difficult to interpret MIC results or inconsistent results across replicates. Some test strips were discontinued during the course of this experiment and were therefore marked as missing in the antibiogram.

Antibiograms were obtained for the majority of isolates across all tested drugs (Table S2) using E tests; selected drugs were chosen based on treatment suggestions in previous publications (58–68). The MIC values were mapped against a phylogeny of *A. baumannii* genomes (Figure 1) inferred from a concatenation of 182,916 SNPs. Resistant, susceptible, or intermediate calls were determined based on identified breakpoints (Table 1). Four of the drugs used in this study do not have an identified breakpoint for *Acinetobacter*. We applied breakpoints for two of these drugs based on other organisms. For two additional drugs, breakpoints were applied that only includes the highest and lowest values of the E test range. This conservative approach is potentially useful for grouping isolates into categories to identify mechanisms associated with the largest differences in MIC values, but may not be clinically or biologically relevant.

From the *A. baumannii* phylogeny, isolates were identified that were closely related based on phylogenetic distance, but differed in their antibiograms. These isolate pairs (Figure 2, Table 2) were the subject of additional investigation in order to identify cryptic resistance mechanisms based on a common genomic background.

**Figure 2:**
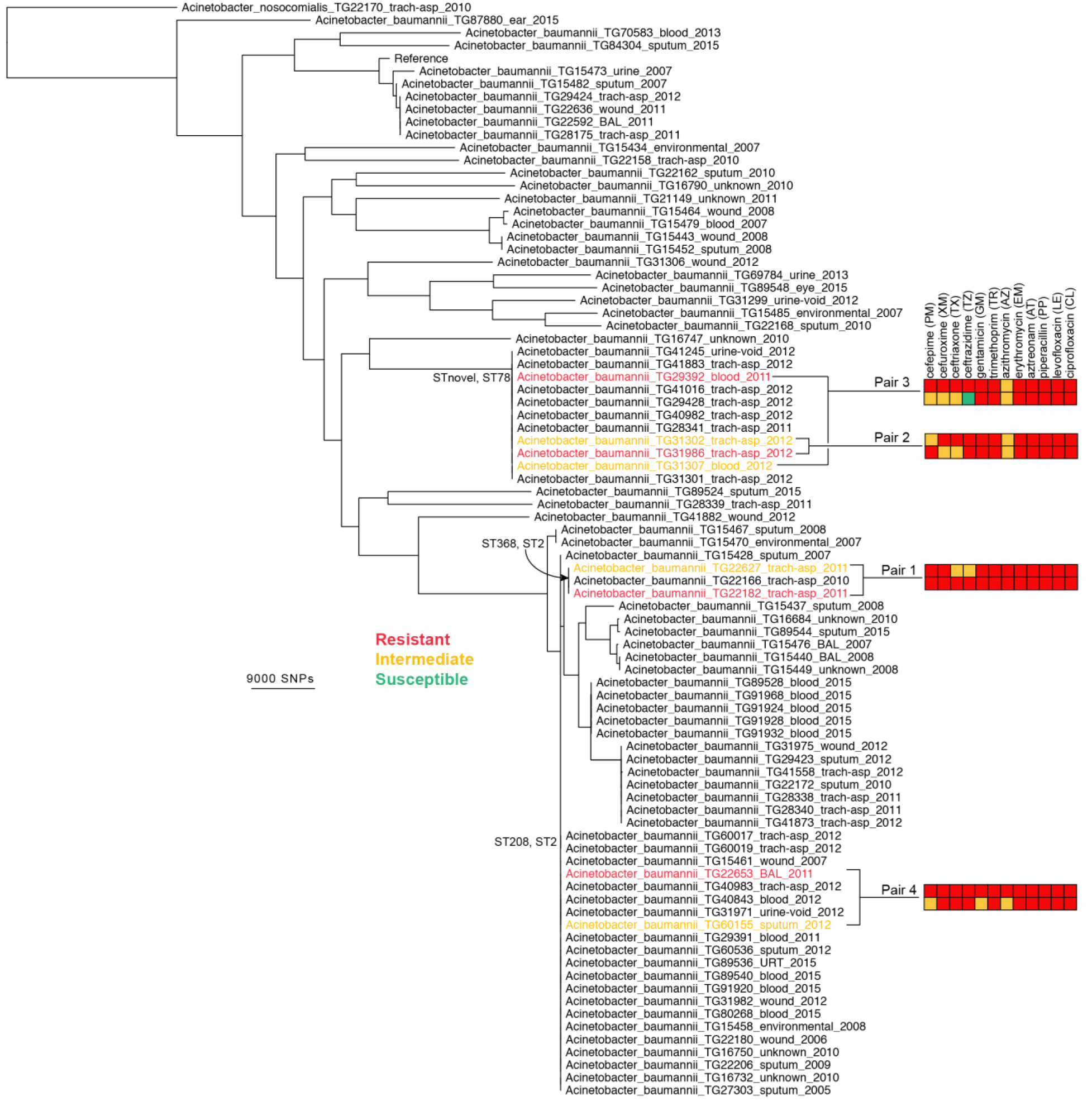
A maximum-likelihood phylogeny of *A. baumannii* genomes sequenced in this study, based on a concatenation of 182,766 core genome SNPs. The resistance profiles are shown across paired genomes.

### *in silico* AMR profiling of all sequenced *Acinetobacter* isolates

All proteins from the CARD database (n=2,420) were aligned against 107 sequenced genomes with LS-BSR in conjunction with Diamond. Proteins that were highly conserved in at least 5 genomes were mapped against the phylogeny and demonstrate variable conservation of AMR-associated proteins (Figure 3, Table S5). Some proteins had a clear phylogenetic distribution, where they were either conserved across almost all *Acinetobacter* (e.g. OXA-64 (OXA-51 family)), conserved across phylogenetic groups in *A. baumannii* (e.g. aminoglycoside resistance genes (APH-6): AAC23556.1), or were variably conserved (e.g. aminoglycoside adenyltransferase (ANT(2”)-Ia): AAC64365.1), suggesting horizontal gene transfer.

**Figure 3:**
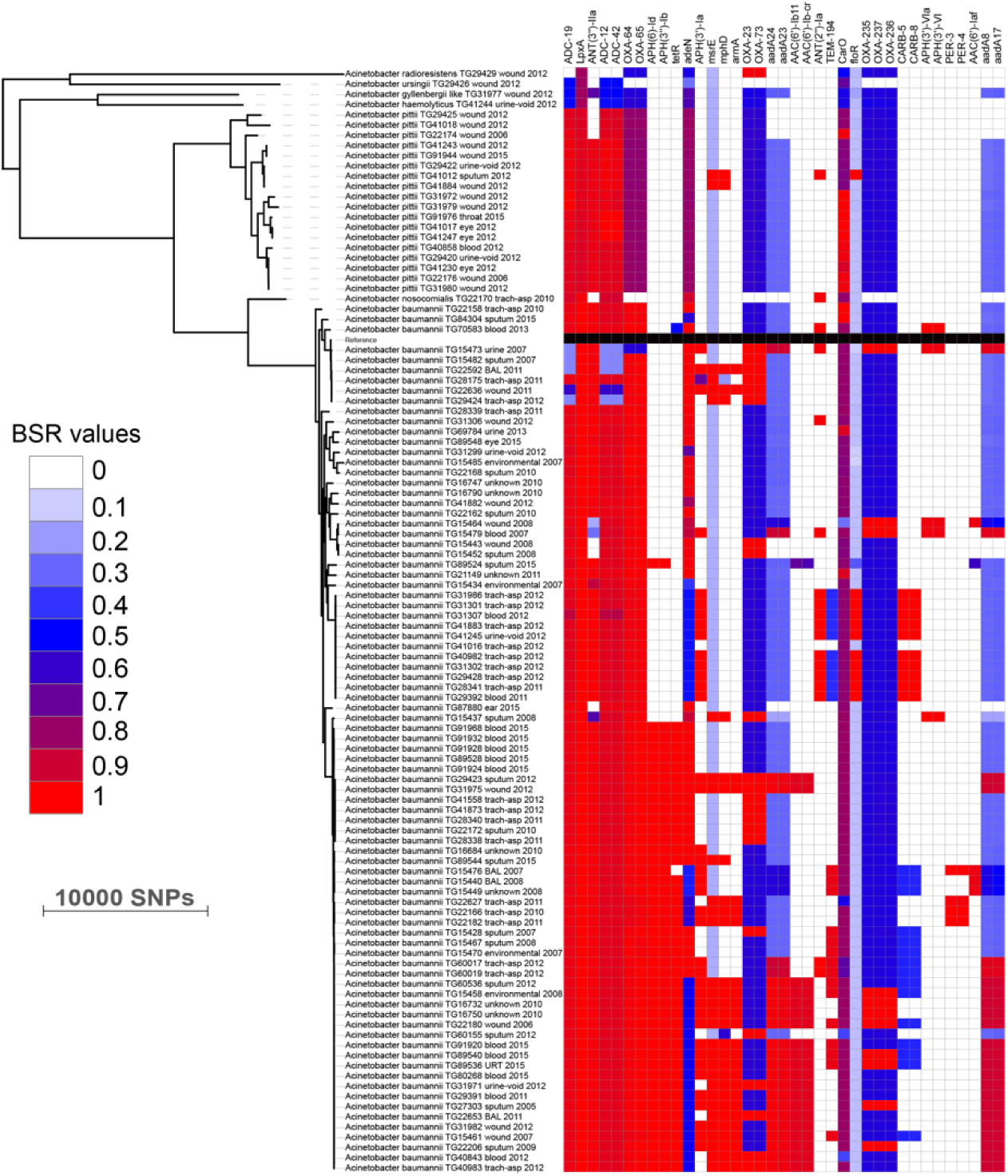
Screen of selected CARD proteins across all *Acinetobacter baumannii* genomes sequenced in this study. The phylogeny is the same as is shown in Figure 1. The heatmap is associated with the blast score ratio (BSR) (77) values of each region across each genome. The BSR values were visualized with the Interactive tree of life (111).

**Table 5:**
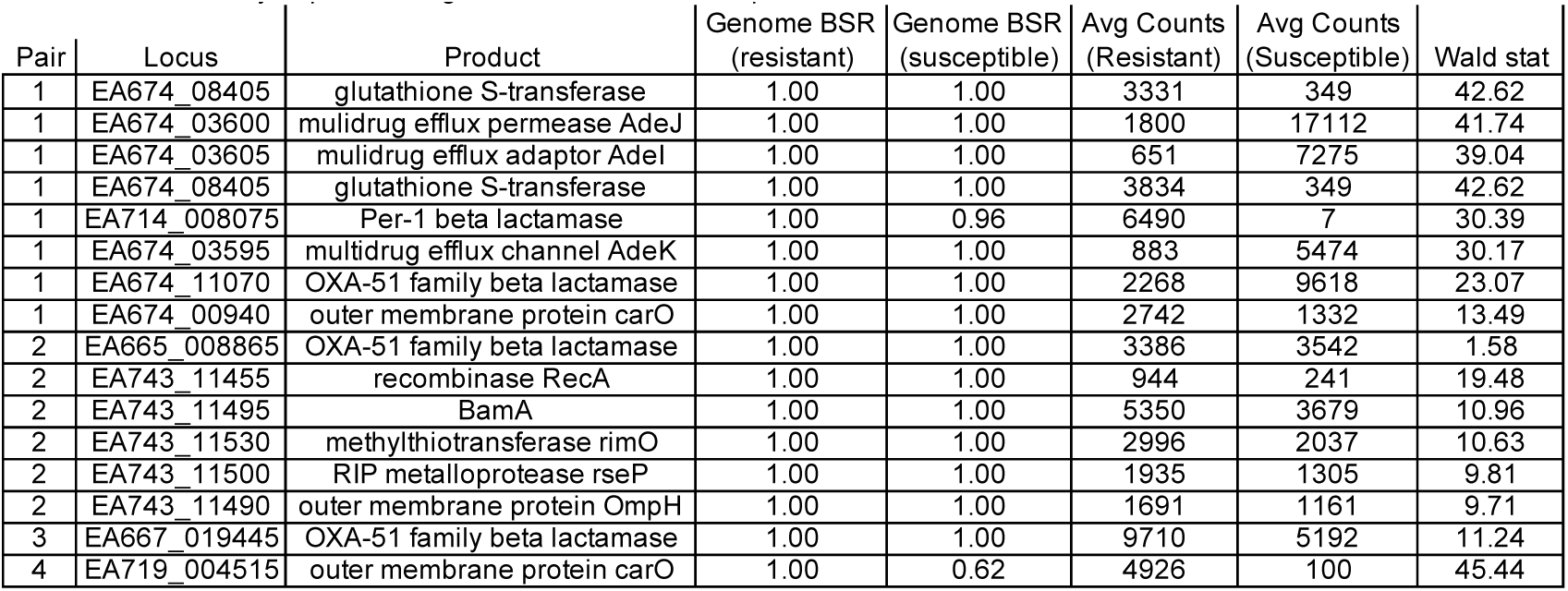
Differentially-expressed regions based on RNA-Seq

| Pair | Locus | Product | Genome BSR<br>(resistant) | Genome BSR<br>(susceptible) | Avg Counts<br>(Resistant) | Avg Counts<br>(Susceptible) | Wald stat |
| --- | --- | --- | --- | --- | --- | --- | --- |
| 1 | EA674_08405 | glutathione S-transferase | 1.00 | 1.00 | 3331 | 349 | 42.62 |
| 1 | EA674_03600 | multidrug efflux permease AdeJ | 1.00 | 1.00 | 1800 | 17112 | 41.74 |
| 1 | EA674_03605 | multidrug efflux adaptor AdeI | 1.00 | 1.00 | 651 | 7275 | 39.04 |
| 1 | EA674_08405 | glutathione S-transferase | 1.00 | 1.00 | 3834 | 349 | 42.62 |
| 1 | EA714_008075 | Per-1 beta lactamase | 1.00 | 0.96 | 6490 | 7 | 30.39 |
| 1 | EA674_03595 | multidrug efflux channel AdeK | 1.00 | 1.00 | 883 | 5474 | 30.17 |
| 1 | EA674_11070 | OXA-51 family beta lactamase | 1.00 | 1.00 | 2268 | 9618 | 23.07 |
| 1 | EA674_00940 | outer membrane protein carO | 1.00 | 1.00 | 2742 | 1332 | 13.49 |
| 2 | EA665_008865 | OXA-51 family beta lactamase | 1.00 | 1.00 | 3386 | 3542 | 1.58 |
| 2 | EA743_11455 | recombinase RecA | 1.00 | 1.00 | 944 | 241 | 19.48 |
| 2 | EA743_11495 | BamA | 1.00 | 1.00 | 5350 | 3679 | 10.96 |
| 2 | EA743_11530 | methylthiotransferase rimO | 1.00 | 1.00 | 2996 | 2037 | 10.63 |
| 2 | EA743_11500 | RIP metalloprotease rseP | 1.00 | 1.00 | 1935 | 1305 | 9.81 |
| 2 | EA743_11490 | outer membrane protein OmpH | 1.00 | 1.00 | 1691 | 1161 | 9.71 |
| 3 | EA667_019445 | OXA-51 family beta lactamase | 1.00 | 1.00 | 9710 | 5192 | 11.24 |
| 4 | EA719_004515 | outer membrane protein carO | 1.00 | 0.62 | 4926 | 100 | 45.44 |

### *in silico* screening of paired isolate genomes

The 84 confirmed *A. baumannii* genomes were screened for the presence of AMR-associated genes from the CARD database with LS-BSR. For 2 isolate pairs, no obvious differences were observed in resistance genes between variably resistant pairs (Figure S2, Table S5). For TG22182 (R) and TG22627 (I), one CARD gene was differentially conserved (CAE51638) and is associated with an aminoglycoside phosphotransferase, although no differences were observed in resistance to the tested aminoglycoside, gentamicin. Multiple differences were observed between the distribution of CARD genes between TG22653 (R) and TG60155 (I) (Figure S2), although the antibiograms only differed in the resistance to two antimicrobials and the genomes differed by only 27 core genome SNPs.

### Screen of previously described AMR mechanisms

A list of mechanisms associated with AMR in *A. baumannii* (Table S3) were screened against genomes sequenced in this study with LS-BSR (Figure S3). Genomes were also screened for mechanisms associated with resistance to the following specific drugs:

#### Quinolones

The *gyrA* S82L mutation has previously been shown to confer resistance to quinolones in *Acinetobacter* (29). Of the resistant *A. baumannii* strains (n=73), 72 (∼99%) contained the leucine (L) residue at position 82; the one exception was TG22162, which had the serine (S) residue. All susceptible strains had the serine residue at this position, suggesting that this mutation is the primary mechanism conferring quinolone resistance in analyzed strains.

#### Trimethoprim

All tested *A. baumannii* isolates were resistant to trimethoprim. An *in silico* screen of *folA* (Figure S3), which has been associated with target site alteration and trimethoprim resistance (25), demonstrated that all genomes contained this gene, although there was some variation in the peptide identities. As susceptible strains were not identified through screening, we cannot test the genotype/phenotype relationship for this compound, although based on published results, *folA* appears to be the associated mechanism.

#### Beta-lactams

The *ampC* gene in *A. baumannii* is a class C beta-lactamase (87). A screen of the *ampC* peptide sequence against *A. baumannii* isolates sequenced in this study indicates that almost all genomes have a highly conserved *ampC* gene at the nucleotide level, but have widely different antibiograms (Table S2, Figure S3). This demonstrates that the presence/absence of this gene alone has little predictive value on beta-lactam resistance in *A. baumannii*.

The insertion element ISAba1, in conjunction with bla_OXA-51-like_ genes, has been shown to confer resistance to carbapenems (22). Genomes in this study showed a correlation (>0.8 correlation coefficient) between ISAba1 conservation and resistance to 2 beta-lactams (XM,TX). Many of the ISAba1 transposases were split across multiple contigs in Illumina assemblies, likely due to a repeat region that could not be resolved during assembly. Furthermore, copies of this region were likely collapsed during the short read assembly. For example, for the 8 isolates for which draft genomes and complete genomes were generated in this study, only a single copy of ISAba1 was observed in the draft genome, while 9-26 copies were observed in complete genomes. Additionally, all of the paired isolates in this study contained ISAba1 and a bla_OXA-51-like_ gene but showed variable antibiograms to at least one beta-lactam, suggesting that the conservation of this region alone did not explain the resistance phenotype.

Some genes associated with efflux (*adeA*, *adeB*) were missing from several genomes (Figure S3) that showed susceptibility to a number of drug families. Some genomes contained these regions but were also susceptible to beta-lactams, suggesting multiple genotypes result in the same resistance or susceptibility phenotype.

#### Macrolides

Although *ermB* has been associated with macrolide resistance in *A. baumannii*, the gene was not detected in any genome sequenced in this study, based on a LS-BSR analysis (Figure S3). The *mefA* gene was also screened, as it has been demonstrated to provide macrolide resistance, but the gene was highly conserved, even in azithromycin susceptible strains (Figure S3). Resistance to macrolides has also been associated with efflux, although differences in efflux cannot be investigated with genomics alone.

#### Aminoglycosides

Four genes associated with aminoglycoside resistance were screened against genomes with LS-BSR. None of the 4 regions (*aacC1*, *aphA6*, *aadA1*, *aadB*) previously associated with aminoglycoside resistance showed any association (correlation coefficient <0.5 0) with resistance to gentamicin in genomes screened in this study (Figure S3).

### Machine learning approaches

For most of the tested drugs, the machine learning method Kover identified regions that were associated with resistance across the *A. baumannii* isolates tested in this study (Table 3), based on all possible 21-mers. In almost all cases, the frequency of these Kmers could not completely distinguish between resistant and susceptible phenotypes, suggesting that the large number of associated Kmers identified by Kover are likely not biologically meaningful.

### Genome wide association study (GWAS)

To identify genotype/phenotype associations, we performed a GWAS analysis by splitting up isolates into resistant/susceptible groups for each drug; for this analysis we ignored isolates with an intermediate phenotype in order to isolate the mechanism. Using CDS conservation, we failed to identify a clear genotype/phenotype relationship across all *Acinetobacter* across all drugs. This suggests that genomic analyses alone can fail to comprehensively identify AMR mechanisms in *Acinetobacter*. When the analysis was repeated for only *A. baumannii* genomes using both coding regions and Kmers, significant associations were identified (Table 4). For ceftriaxone (TX), all susceptible strains (n=3) were missing ISAba1 (ABLAC_32600), while 66 of 67 resistant strains contained this region; the small number of genomes analyzed limits the power of this analysis. In spite of this correlation, the lack of broadly conserved genomic regions associated with resistance directed a paired genome analysis into the identification of novel or cryptic genotype/phenotype associations.

### RNA-Seq and differential expression (DE) analysis

For isolate pairs where a clear genotype/phenotype relationship was not identified, RNA was extracted and cDNA was sequenced. Despite implementing methods to enrich mRNA, ∼20% rRNA+tRNA presence was observed in all samples (Table S8). For each pair, all coding and intergenic sequences were combined into a single file and de-replicated. Reads were mapped against these regions, normalized, and differential expression was identified using DESeq2. Results were then identified for the following isolate pairs:

#### Pair 1

Multiple differentially expressed genes were identified that were upregulated in the resistant strain (TG22182) (Table 5). One of these regions was a PER-1 beta-lactamase gene (bla_PER-1_)(EA714_008075) that is not broadly conserved across ST368 genomes (Figure S3) and appears to be present on a transposon. A screen of this gene against other ST368 genomes isolated from diverse geographic locations suggested that there was an acquisition of this region in a single sequence type and a clear phylogenetic effect (Figure S4). Indeed, a genomic island was identified in both the resistant and intermediate genomes between two transposases (Figure 4a) that includes a Glutathione S-transferase gene (EA674_08405) that was also upregulated in the resistant strain (11.2x up-regulation); this region has previously been associated with beta-lactam resistance (88). The operon structure was similar between resistant and intermediate strains, with the exception of an IS91 transposon that was between an IS26 transposon and bla_PER-1_. The operon structure for the resistant strain that contained an IS91 transposon was determined to be highly similar with an ISCR1 (Insertion sequence Common Region) element. Within the ISCR1 element is an ori*IS* (origin of replication) region that allows for rolling-circle replication and transposition of the ISCR1 element. Within the ori*IS* are two outward-oriented promoters (P_OUT_) that have been shown to affect downstream gene expression (89). The resistant strain, TG22182, has both P_OUT_ promoters associated with increased gene expression directly upstream of the bla_PER-1_ gene (Figure 4a). The more susceptible strain, TG22627, has neither of the P_OUT_ promoters upstream of its bla_PER-1_ gene. Additionally, the composition of the bla_PER-1_ gene between isolates was different, with a different coding length as well as composition in the first 12 amino acids of the peptide.

**Figure 4:**
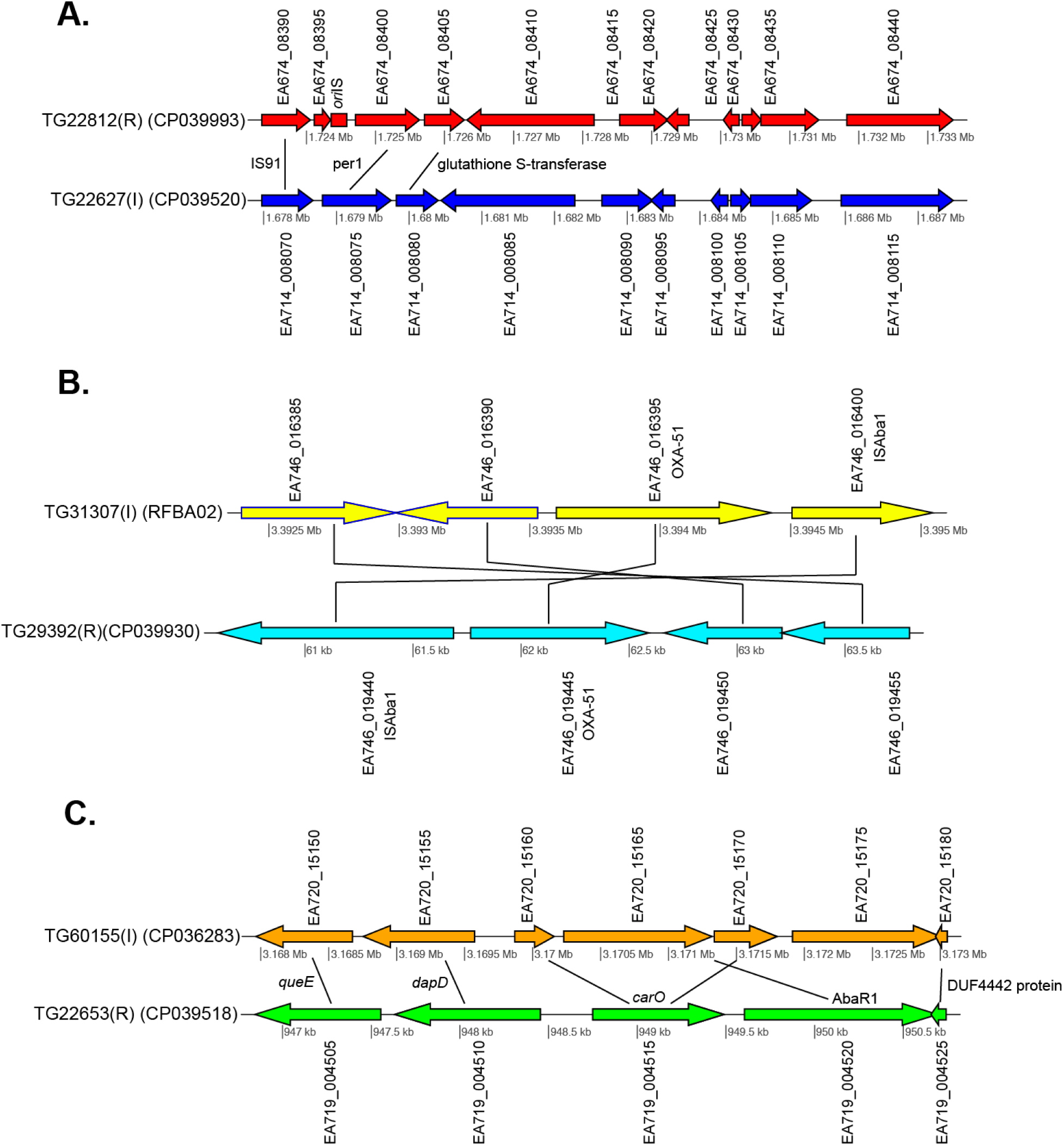
Gene content comparisons between paired isolates in pair 1 (A), pair 3 (B), and pair 4 (C). All figures were generated with genoPlotR (76).

Additionally, a glutathione S-transferase family gene (EA674_008405) and the carbapenem susceptibility porin *carO* (EA674_000940) gene were highly expressed in TG22182 (R) in comparison to TG22627 (I). Both of these genes have been shown to confer resistance to beta-lactams and specifically carbapenems (90). Both isolates in Pair 1 also contain an OXA-51 beta-lactamase gene (bla_OXA-51_) (EA674_011070), although the gene is slightly up-regulated (4.5x) in TG22627 (I). TG22627 also showed higher expression of other genes associated with AMR including *adeI* (EA674_003605)(11x), *adeB* (EA674_009735)(9.5x), *adeA* (EA674_009730)(9.6x), and *adeJ* (EA674_003600)(9.6x); however, the up-regulation of genes in the AdeABC pump have previously been demonstrated to not confer aminoglycoside resistance (23). This suggests that the action of the PER-1 beta-lactamase is more effective than other proteins associated with efflux or other oxacillinases in resistance to ceftriaxone (TX) and ceftazidime (TZ). The PER-1 beta-lactamase protein has also been associated with virulence in *A. baumannii* (91) and has also been associated with resistance to beta-lactams (92). The detection of the bla_OXA-51_ and ISAba1 regions in these genomes revealed nothing about their susceptibility to TX or TZ.

#### Pair 2

Twenty-four differentially-expressed regions were observed between Pair 2 isolates (TG31302, TG31986) at a Wald stat value of 10 (Table S6). None of these regions were associated with known mechanisms of AMR in *A. baumannii*. The two isolates contained a class-D beta-lactamase gene (bla_OXA-51_) and a class-C beta-lactamase gene (*ampC*). There was no significant difference in gene expression of these regions between isolate pairs.

**Table 6:** AmpSeq results

| Pair | Locus | Read<br>counts (R) | Read<br>counts (I) | Control<br>counts (R) | Control<br>counts (I) | AVG<br>delta (R) | AVG.<br>delta (I) | delta-delta | p-value |
| --- | --- | --- | --- | --- | --- | --- | --- | --- | --- |
| 1 | PER-1 (EA714_008075) | 30867 | 309 | 30007 | 62187 | 1549 | 61878 | 60329 | <0.0001 |
| 1 | aphA1 (EA674_13195) | 9172 | 9 | 30007 | 62187 | 23832 | 62178 | 38346 | <0.0001 |
| 2 | *ampC* (EA743_05675) | 48550 | 45017 | 10047 | 10783 | 39002 | 34233 | 4769 | 0.240 |
| 3 | OXA_65 (EA746_016395) | 22818 | 24152 | 30446 | 40298 | 7078 | 16146 | 9068 | 0.0003 |
| 4 | *carO* (EA719_004515) | 31632 | 25088 | 7067 | 30654 | 20926 | 5566 | 15360 | <0.0001 |
| 4 | CsuA/B (EA719_006180) | 23920 | 4 | 7067 | 30654 | 15309 | 30650 | 15341 | <0.0001 |

The composition of each beta-lactamase was determined at the nucleotide and peptide level. A protein alignment of *ampC* between TG31302 (I) and TG31986 (R) revealed a single amino acid difference (R to G) at position 172 (Figure S5) in the PAZ domain. This non-synonymous mutation falls within the second of three characteristic conserved motifs, RxY^150^xN, for all class C serine beta-lactamase sequences (93). Although this mutation has not been previously associated with increased activity or misfolding of the protein, other mutations in *ampC* gene have (94), suggesting that this mutation may confer increased hydrolyzing activity against beta-lactams, although additional validation work is required to test this hypothesis.

A continuous stretch of 17 genes was upregulated in TG31986 (R) (EA743_011445 - EA743_011530) (Table S7). Nine genes within this region were in the top 30 differentially expressed genes in this analysis, including: recombinase *recA* (EA743_011455), outer membrane protein assembly factor *bamA* (EA743_011495), 30S ribosomal protein S12 methylthiotransferase *rimO* (EA743_011530), UMP kinase gene *pyrH* (EA665_011490), RIP metalloprotease *rseP* (EA743_011500), a gene coding for an OmpH family outer membrane protein (EA743_011490), a phosphatidate cytidylyltransferase gene (EA665_011475), a 1-deoxy-D-xylulose-5phosphate reductoisomerase gene (EA665_011470), and a di-trans,poly-cis-decaprenylcistransferase gene (EA665_011480). The outer membrane protein (OMP) H, a homolog of the Skp protein in *E. coli* (95), has been classified as a chaperone protein involved in the folding of BamA. Previous research has shown a correlation between upregulation of molecular chaperones when exposed to antimicrobials and the bacterium’s improved ability to tolerate antimicrobial stress (96). Researchers have demonstrated that the *skp* gene in *E. coli* is an important stress-associated gene (97) that may be associated with AMR (98).

#### Pair 3

Of 94 genomic regions that were significantly differentially expressed (Walt stat >10 or <-10) (Table S8) between this isolate pair, one of the significant differences was between an bla_OXA-51_ family (OXA-65) beta-lactamase gene (EA667_019445), which showed 1.9x up-regulation in the resistant strain. Interestingly, 79 bases separated the end of insertion element ISAba1 and the start codon of bla_OXA-51_ in sample TG29392 (R). Previous research has demonstrated that this bla_OXA-51-like_ gene is conserved across *A. baumannii* lineages, but only genomes containing the ISAba1 directly upstream of bla_OXA-51_ show resistance to carbapenems. It is likely that ISAba1 is acting as the promoter for bla_OXA-51_ in TG29392, conferring resistance to carbapenems (22). The intermediate resistance strain, TG31307, has both ISAba1 and bla_OXA-51_; however, ISAba1 is downstream of bla_OXA-51_ and therefore not functioning as a strong promoter for the beta-lactamase gene (Figure 4b). An analysis of the coding region of the bla_OXA-51_ gene in both isolates revealed no differences, suggesting that differences in expression are due to expression.

Interestingly, several of the CDSs that were differentially expressed in the resistant strain in Pair 2 were up-regulated in the intermediate Pair 3 strain (TG31307). For example, *recA* was the most differentially expressed gene in Pair 3 genomes, but was up-regulated in TG31307 (Table S8). This suggests that the mechanisms of resistance have complex interactions that need to be investigated through targeted gene deletions.

#### Pair 4

Of the numerous differences in gene expression between the resistant (TG22653) and intermediate strain (TG60155), perhaps the most striking is in the expression of *carO* (EA719_004515), an outer membrane porin (Table S9). Previous analyses have demonstrated that insertion sequences that disrupt *carO* are associated with decreased activity against beta-lactams (99). The genome assembly of the intermediate strain, TG60155, shows that *carO* is interrupted by the insertion sequence, ISAba1 (EA720_015165) (Figure 4c). An analysis of the transcriptome of TG60155 also failed to identify an intact *carO* transcript, which was present in TG22653.

### Antimicrobial resistance induction analysis

In an effort to observe induced antimicrobial resistance in the four paired isolates, resistant strains were grown in sub-inhibitory concentrations of select antimicrobials. Differential expression of each sub-inhibitory isolate was compared to the isolate grown under inhibitory concentrations using the Wald statistic produced from DESeq2. Additionally, the resistant isolate TG22653 was grown under two different sub-inhibitory concentrations of antimicrobials (16 and 258); differential expression from these two concentrations was also compared. No significant differential expression was observed in the four analyses based on an FDR-adjusted p-value of 0.05. Likewise, using the Wald statistic from these analyses also demonstrated no significant differential expression between the differing antimicrobial concentrations using the chosen threshold. This suggests that differential expression is due to constitutively expressed mechanisms that are not inducible.

### AmpSeq validation

Amplicon sequencing was performed on cDNA to not only confirm the RNA-Seq results, but also to provide a proof of concept as an advanced AMR diagnostic. Comparative expression was identified through comparison of ratios of the number of read counts of each targeted gene compared to a housekeeping gene (IX87_18340); the housekeeping gene was identified as a gene with consistent, and relatively high, expression in the RNA-Seq data. For pair 1, the bla_PER-1_ and aphA1 genes were significantly upregulated in the resistant strain compared to the intermediate strain (Table 6). For pair 2, the expression of the *ampC* gene was not significantly different, suggesting that differential expression of this region doesn’t explain the resistance phenotype and is consistent with the RNA-Seq data. For pair 3, the bla_OXA-51_ gene was confirmed to be significantly up-regulated in the resistant strain compared to the intermediate strain. For pair 4, the *carO* gene was significantly up-regulated in the resistant strain, which is consistent with the RNA-Seq results and is likely the primary mechanism of resistance. A gene associated with the production of a spore-coat forming protein (CsuA/B) was highly up-regulated in the resistant strain. While not directly associated with AMR, this validation provides confidence in the RNA-Seq data.

### General transcriptome screen

A LS-BSR analysis of previously described resistance mechanisms (Table S3) between the genome and transcriptome demonstrated that some genomic regions, such as the *adeF* gene, were highly conserved in the genome, but were largely absent from the transcriptome (Table S10). This finding demonstrates the importance of incorporating gene expression when trying to understand phenotypic differences.

## Discussion

Antimicrobial resistance (AMR) is a significant, emerging threat, with *A. baumannii* being recently classified as a priority 1 pathogen (2). Some mechanisms associated with AMR in *A. baumannii* are clearly understood, especially with regards to documented beta lactamases (100–103) and efflux pumps (15, 104, 105). However, the highly plastic pan-genome of *A. baumannii* (45) suggests that the identification of universal AMR mechanisms may be unlikely, even with regard to the presence and activity of specific beta-lactamases. This same trend has been observed in other highly plastic genomes, such as *Pseudomonas aeruginosa* (106), and complicates surveillance and targeted therapy efforts. As such, *A. baumannii* is not only an emerging threat, but represents a critical challenge to the development of both novel drugs and molecular diagnostics.

In this study, we sequenced 107 genomes reported to be *A. baumannii* based on testing in the clinical laboratory. Typing based on WGS analyses identified 23 of the genomes were misclassified and belonged to other *Acinetobacter* species (Table S1). These incorrect clinical laboratory typing results highlight the need for improved clinical diagnostics of *A. baumannii*. An additional 35 genomes in the GenBank assembly database were incorrectly annotated as *A. baumannii* and belonged to other species (not shown), which further demonstrates the difficulty in typing as well as and the impact of mis-annotated genomes on population structure analysis in *Acinetobacter*. Typing strains using WGS should be a first step in any large comparative genomics study to limit the analysis to a targeted group, clade, or species.

We generated antibiograms for 12 drugs, representing multiple drug families, across the 84 isolates confirmed to belong to *A. baumannii* by WGS analysis. Some of the drugs screened in this study aren’t typically used in current treatment regimens for *A. baumannii* infections. However, with growing resistance emerging to next generation drugs, clinicians are exploring older drugs (e.g. chloramphenicol) to treat emerging threats (42, 107). In this study, we sought to identify genomic differences that could explain the variable resistance phenotypes using established antimicrobial resistance gene databases as a method to predict AMR from genomics data (31, 32, 35, 60). A screen of regions from the Comprehensive Antimicrobial Resistance Database (CARD) against genomes sequenced in this study failed to identify characterized resistance mechanisms that largely explain the resistance phenotype. These results demonstrate the limitations to this approach in highly plastic species, such as *A. baumannii*, and suggest that alternative approaches including RNA-seq data may be required for a comprehensive understanding of AMR mechanisms in *A. baumannii*.

We then employed reference independent, genome wide association study (GWAS) methods to identify genomic differences between susceptible/intermediate/resistant phenotypes. These types of associations have been used in other pathogens to identify genotype/phenotype associations (108). In general, we failed to identify a clear association between the genotype (21bp Kmers, coding regions, SNPs) and the resistance phenotype when comparing either all *Acinetobacter* genomes or just *A. baumannii* genomes (Table 4). This result suggests that diverse and independent mechanisms may be responsible for the AMR phenotype for some drugs instead of single, highly conserved mechanisms.

Recent research has demonstrated difficulty identifying complex mechanisms, or under-or-over-represented phenotypes, using a GWAS approach (109). As a way to focus on sparsely distributed AMR mechanisms, a paired isolate approach was utilized in order to reduce noise in the genomic background. In this analysis, four isolate pairs were individually compared across four antimicrobials (Table 2). RNA-Sequencing (RNA-Seq) of these four paired resistant/intermediate isolates that shared a common genetic background was employed. Using this approach, several potential mechanisms were identified, some of which have been identified previously in *A. baumannii*, but are difficult to identify with standard comparative genomics approaches, requiring RNASeq for comprehensive surveillance. While we identified known resistance mechanisms in the resistant strains, some of those regions were also identified in intermediate strains; previous studies of *A. baumannii* transcriptomes have also observed up-regulation of resistance and efflux genes in susceptible strains (5). RNASeq data allowed for the identification of antimicrobial resistance mechanisms that while present in both the resistant and susceptible genomes, were differentially expressed due to an upstream insertion element. These results also highlight the possibility that expression of a single AMR gene does not always confer resistance and it is likely that combinations of genes are responsible for observed resistance. Differential expression differences were confirmed using a cDNA-AmpSeq approach and largely confirmed the differential expression of targeted regions. Previous research has demonstrated a bias of differentially expressed regions when applying a multiplexed PCR approach (110). We addressed this issue by optimizing primer concentrations using genomic DNA and including a single copy number gene for normalization.

The results of this study demonstrate that, due to the plastic nature of *A. baumannii’s* pan-genome, comprehensive AMR surveillance cannot solely be achieved through genomics methods alone, especially with current AMR databases and commonly used analytical methods. This study demonstrates that AMR genes are not conserved across *A. baumannii* lineages with similar AMR profiles and that solely relying on genomics methods for AMR surveillance and discovery, such as gene presence/absence, will fail to detect novel or recently acquired AMR mechanisms. For instance, identifying only the position of insertion sequence (IS) elements throughout a genome using genomic tools provides little resolution to inform of possible AMR genes upregulated by the presence of upstream IS elements. Furthermore, identification of these elements would provide little resolution of antimicrobial resistance profiles in a clinical setting. However, by utilizing transcriptome data, we were able to identify AMR genes upregulated by these elements as well as novel AMR mechanisms, and design rapid cDNA amplicon sequencing targets for these mechanisms to improve surveillance and diagnostic efforts.

---

**Figure S1:**
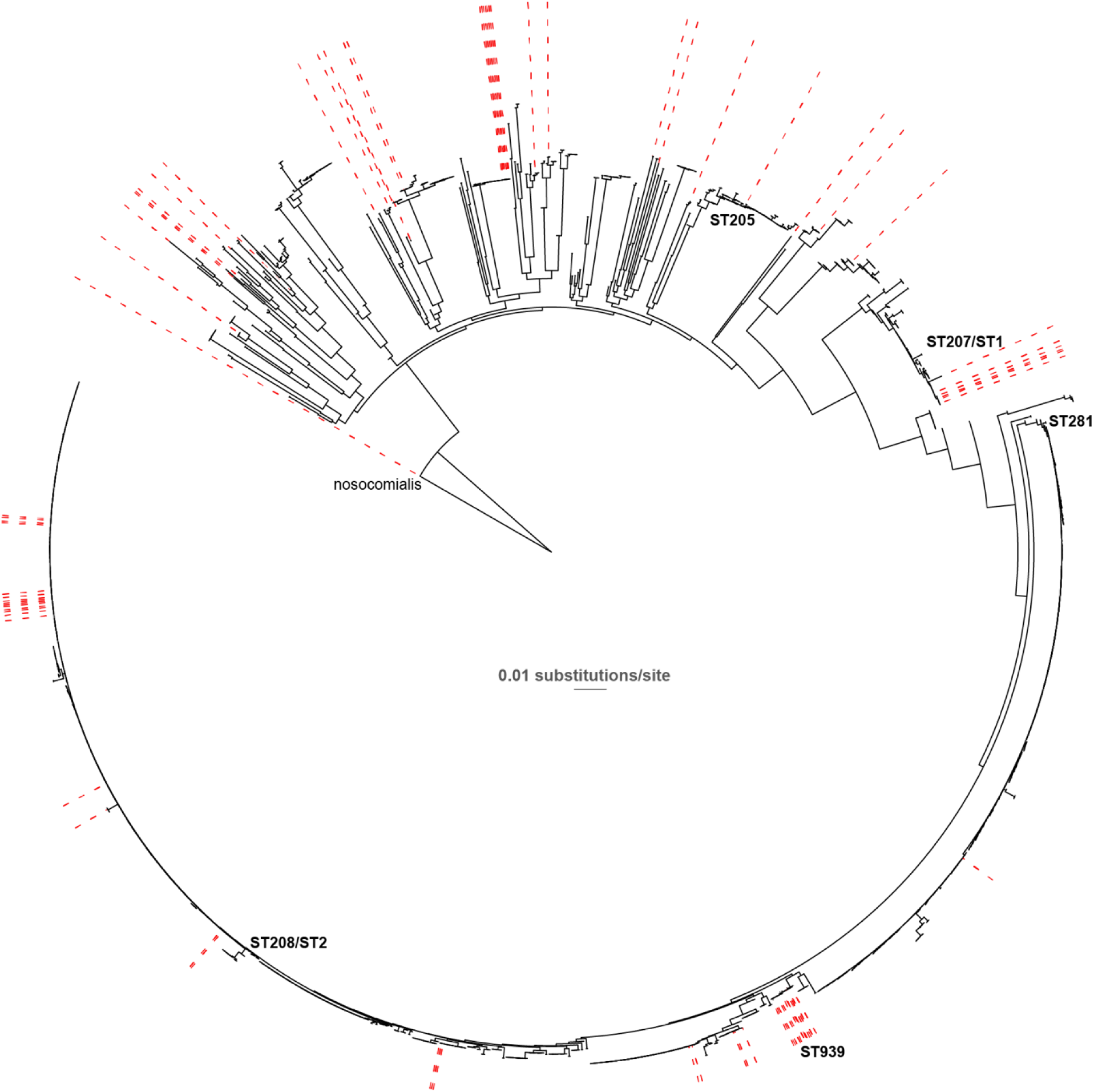
A maximum-likelihood phylogeny of global *A. baumannii* genomes inferred from an alignment of 11,687 concatenated SNPs. Red dashes point to genomes sequenced in this study.

**Figure S2:**
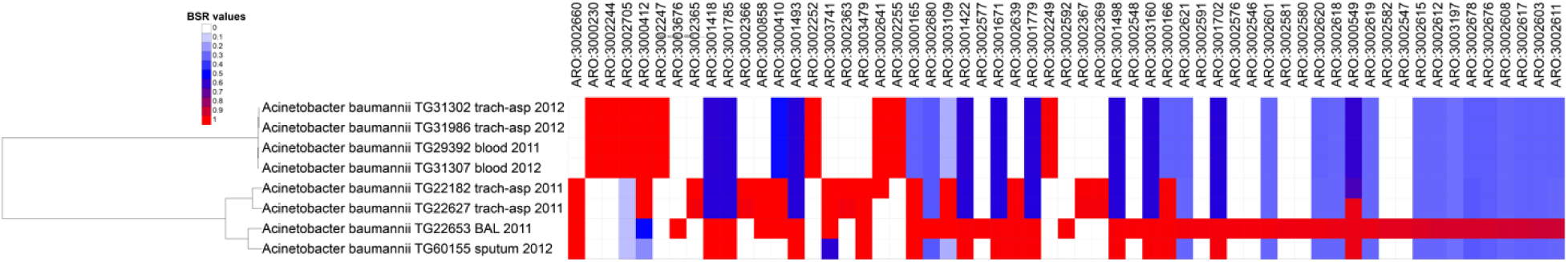
A maximum-likelihood phylogeny of paired isolates. The conservation of selected proteins from the CARD database, based on blast score ratio (BSR) values, is shown as a heatmap. The BSR values were visualized with the Interactive tree of life (111).

**Figure S3:**
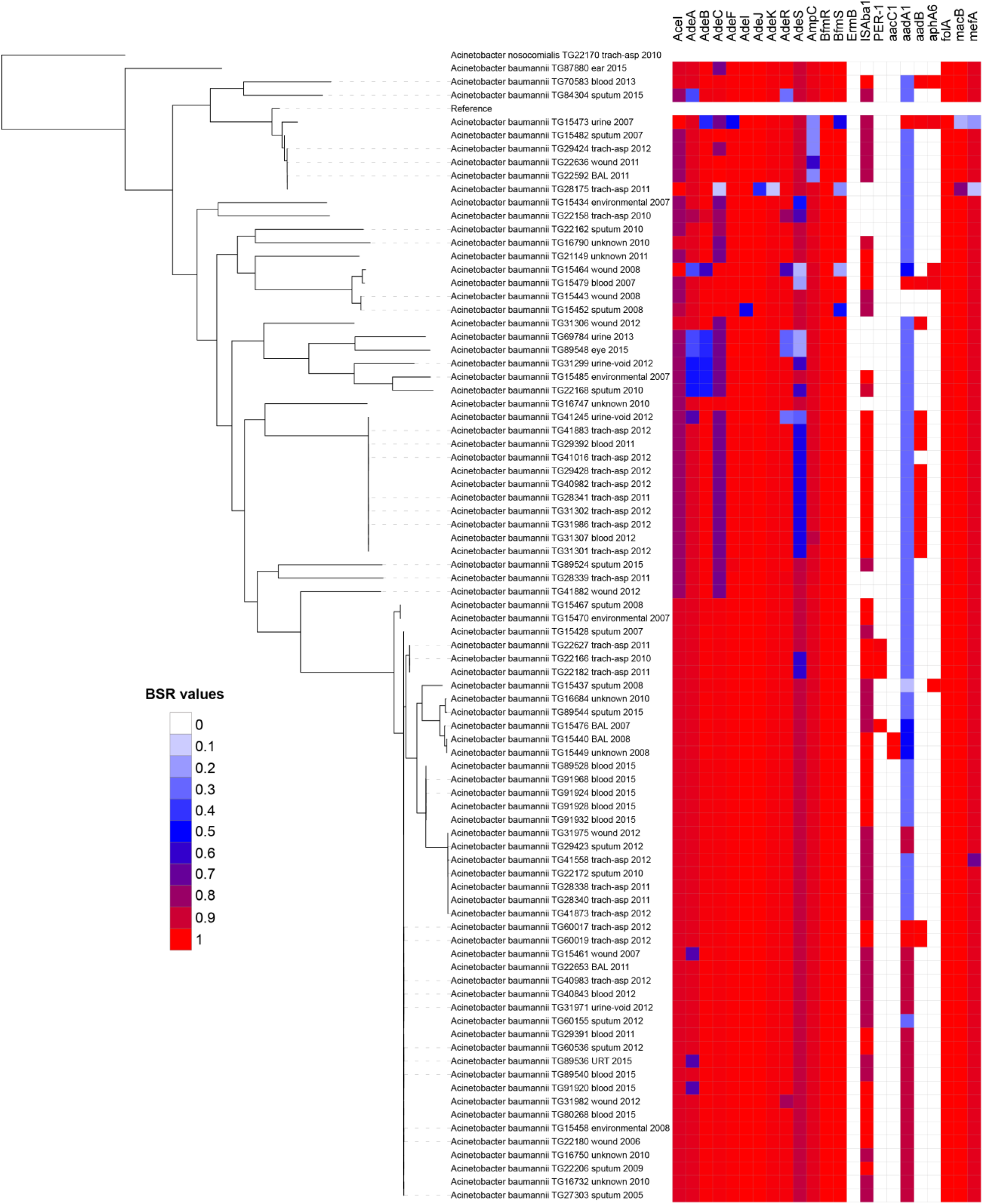
A maximum likelihood phylogeny of *A. baumannii* genomes. The blast score ratio (BSR) of genes associated with AMR (Table S3) were visualized as a heatmap with the Interactive tree of life (111).

**Figure S4:**
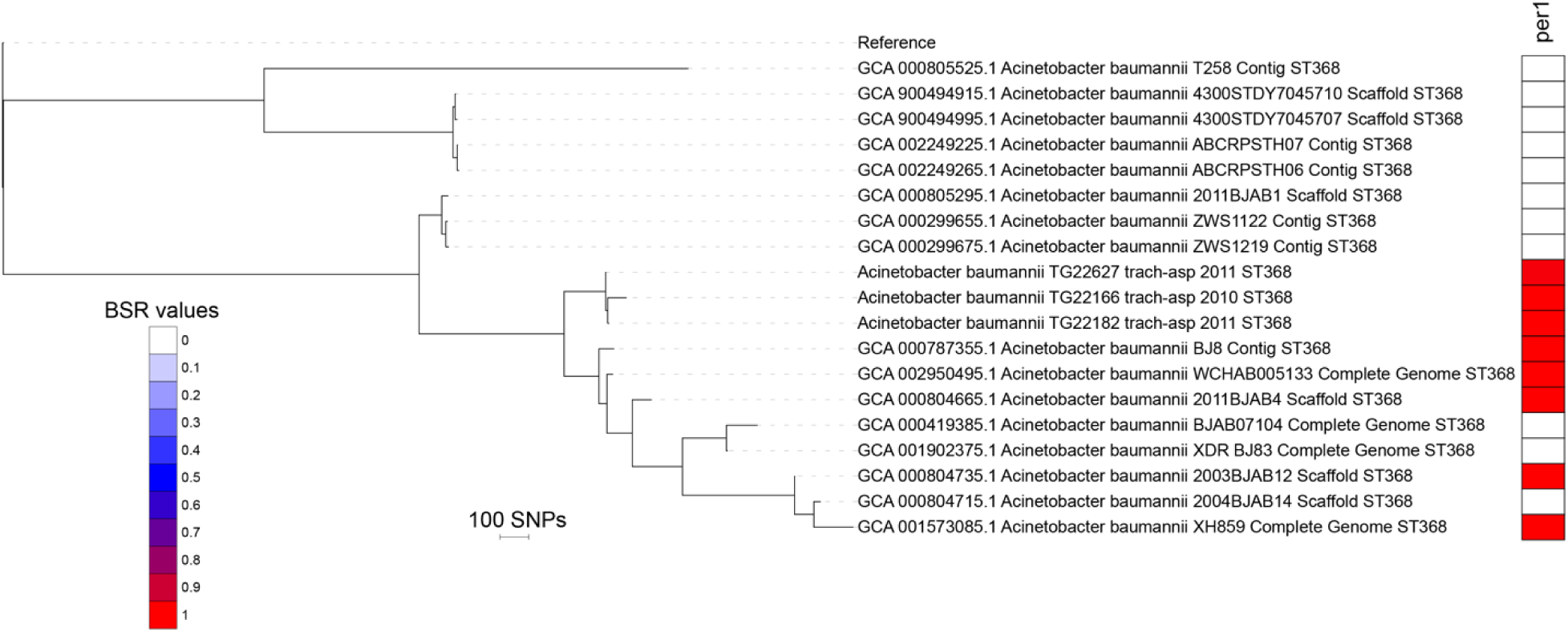
A maximum likelihood phylogeny of select *A. baumannii* genomes. The distribution of the bla_PER-1_ beta-lactamase gene in ST368 genomes, based on blast score ratio (BSR) values, was visualized as a heatmap with the Interactive tree of life (111).

**Figure S5:**
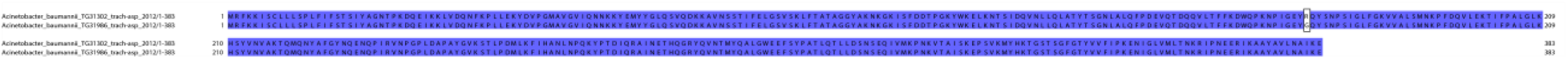
A peptide alignment of *ampC* from pair 3 genomes. The variable residue is outlined with a black box. The alignment was visualized with JalView (112).

